# Willingness of Japanese people in their 20s, 30s and 40s to pay for genetically modified foods

**DOI:** 10.1101/2023.10.29.564581

**Authors:** Akihiro Mine, Sawako Okamoto, Tomoya Myojin, Miki Hamada, Tomoaki Imamura

**Affiliations:** Department of Public Health, Health Management and Policy, Nara Medical University, Kashihara, Nara, Japan; Advanced Service Development Group, Innovation Service Creation Division, Mitsubishi Research Institute, Inc., Tokyo, Japan

## Abstract

The application of genetically modified (GM) technology to food products has increased worldwide. The adaptation has extended to conventional grains and animal products, such as salmon. However, in Japan, the public’s acceptance of GM foods is low and experts and policymakers need to know the public’s preference for various types of GM foods. Therefore, this study aims to clarify and compare the preferences for various GM foods among Japanese people in their 20s, 30s, and 40s, using the Willingness-to-pay (WTP) indicator. An online survey with 1122 valid responses from people in their 20s-40s was used for analysis. The results showed that the percentages of willingness to purchase various items were as follows – GM blue roses (n = 628; 56%), tomatoes fertilized with GM oil cake (n = 519; 46.3%), potato chips made from GM potatoes (n = 489; 43.6%), chicken thighs fed on GM corn (n = 472; 42.1%), corn flakes made from GM corn (n = 471; 42.0%), GM apples (n = 420; 37.4%), wine brewed with GM yeast (n = 416; 37.1%), GM tomatoes (n = 408; 36.4%), GM chicken thighs (n = 360; 32.1%), GM salmon (n = 349; 31.1%). Comparing the WTP discount rates for animal and plant foods, it was around 20% for animal products (GM chicken thighs = 17.8% and GM salmon = 19.5%), and 15–35% for plant foods (GM corn flakes = 24.1%, GM tomatoes = 17.2%, GM potato chips = 16.4%, GM yeast wine = 36.7%, GM apples = 25.3%). The WTP discount rates were 11.9% for tomatoes fertilized with GM oil cake compared with 17.2% for GM tomatoes, and 21.5% for GM-fed animal products compared with 17.8% for chicken thighs fed with GM corn. Therefore, the WTP values of GM animal foods were lower than GM plant foods and ornamental products.

## Introduction

Various types of genetically modified (GM) food have been developed and introduced in the market in the last few decades [1]. However, investigations have shown that the public acceptance of GM food has not increased worldwide, especially in European countries and Japan [2–7].

GM techniques are applied to vegetables and animals [1]; therefore, it is imminent that these GM foods will come on the market. In Japan, despite people’s low acceptance, the Japanese government has approved several import crops for food, feed, and processing [1,8]. Since individual preferences for GM foods may differ based on applications [9], it is important to know people’s acceptance of various types of GM foods and their willingness- to-pay (WTP) for GM foods in the market.

The contingent valuation method (CVM) is widely used to quantify individual WTP in the health and environment fields [10]. The WTP allows the transfer of individual preference to a monetary value to compare the preference of individual or varied items [11]. Regarding GM foods, several examinations have been carried out to clarify the acceptance of various GM foods in different regions [12].

This study aims to clarify and compare the WTP of Japanese people in their 20s-40s for various GM products, such as vegetables, animals, feed foods, and ornamental roses.

## Materials and methods

### Data collection

The data used in this study was based on the study that examined the effect of basic biology education in high school on Japanese people in their 20s, 30s, and 40s [7]. The online questionnaire survey was conducted from March to April 2016 by Survey Research Center Co., Ltd., a private research company with a database of Japanese consumers. The company implemented a cutoff of 400 for enrollment in each age group. Finally, 1594 invitation e-mails were sent and 1122 valid responses were obtained, excluding the respondents who sent incomplete responses and those who worked in agriculture-related areas, such as farmers, fishermen, and researchers.

The survey items included sociodemographic characteristics and the WTP of various GM products. The WTP data reported in this study has not been published previously. Regarding sociodemographic characteristics, we gathered information about sex, age, household income (divided into 11 groups by ¥1 million [US$1 = ¥146.5 as of August 28, 2023]), academic background, residence, and whether they had children and the age groups of their children (below 6 years, 6–11 years, and 12–19 years). If respondents had more than one child, the youngest child was considered.

The participants were asked about their WTP in two-step. First, they were asked whether they were “willing” or “unwilling” to purchase 10 GM-related products (cornflakes made from GM corn, tomatoes fertilized with GM oil cake, GM tomatoes, potato chips made from GM potatoes, chicken thighs fed on GM corn, GM chicken thighs, wine brewed with GM yeast, GM apples, GM salmon and GM blue roses). Second, respondents who indicated their WTP were asked how much they were willing to pay for each product. Once the responses were collected, the average market price of corresponding non-GM products at the time was provided for comparison.

### Data analysis

We compared the WTP percentage for the 10 GM products. Furthermore, we calculated the average WTP by using two methods. First, the percentage drop in WTP from the market price of the non-GM product was calculated for those who indicated the WTP. Second, as a sensitivity analysis, the WTP for those without purchase intention was set to 0, and the average was calculated (adjusted WTP). Based on this, the rate of decline from the market price of non-GM alternatives was calculated.

We examined the effect of sociodemographic characteristics, such as sex, age, income, final education level, and the presence or absence of children, on willingness to purchase and WTP. We statistically tested the difference in purchase intention (two values – may or may not purchase) with the χ^2^ test. In addition, we tested whether there was a difference in the WTP, using a t-test for sex and place of residence, and a one-way ANOVA for age, education, income, and presence of children.

This study was approved by the Ethics Committee of Nara Medical University (authorization code: 655). Before answering the questionnaire, the participants provided written consent. They were informed that their responses would be made anonymous and used only for research purposes, and they could withdraw their consent at any time while answering the questionnaire; however, not after submitting their responses. Only when they agreed to these instructions did they click the button to answer the questionnaire.

This research was funded by a Health and Labor Sciences Research Grant (#H27- FoodGeneral-004) in 2016. However, the funding source had no role in the study design, data collection, data analysis, and the decision to submit the manuscript for publication.

## Results

### Sociodemographic characteristics

Table 1 shows participants’ characteristics. Online questionnaires were distributed to 1594 people and 1122 respondents (70.4%) provided valid responses. About half of the respondents were male (46.9%). The percentage of people in their 20s, 30s, and 40s were 31.5%, 32.8%, and 35.7%, respectively. The median of higher income was ¥5-6 million, slightly higher than that of the average Japanese population. Most participants had university liberal arts education (36.6%), followed by vocational school (20.3%), and a small number with postgraduate education (2.9% and 5.3% for liberal arts and sciences, respectively). The majority of respondents had no children living with them (69.1%), followed by those with children below 6 years, 6–11 years, and 12–19 years of age (18%, 7.6%, and 5.3%, respectively).

**Table 1.** Sociodemographic characteristics of enrolled participants (n = 1122).

| Variable |  | n | % |
| --- | --- | --- | --- |
| Sex | Male | 526 | 46.9 |
|  | Female | 596 | 53.1 |
| Age | 20s | 354 | 31.5 |
|  | 30s | 368 | 32.8 |
|  | 40s | 400 | 35.7 |
| Income | Median household income | ¥500–600 million |  |
| Education | High school degree | 170 | 15.2 |
|  | Higher education institution | 228 | 20.3 |
|  | University degree (humanities) | 411 | 36.6 |
|  | University degree (science) | 221 | 19.7 |
|  | Graduate degree (humanities) | 33 | 2.9 |
|  | Graduate degree (science) | 59 | 5.3 |
| Children | No children in the household | 775 | 69.1 |
|  | Below 6 years | 202 | 18 |
|  | 6–11 years | 85 | 7.6 |
|  | 12–19 years | 60 | 5.3 |
| Residence | Rural | 454 | 40.5 |
|  | Urban | 668 | 59.5 |

### Willingness to purchase for GM items

Table 2 shows the percentage of willingness to purchase for various GM items. In descending order of willingness to purchase, GM blue roses (n=628, 56.0%), tomatoes fertilized with GM oil cake (n=519, 46.3%), potato chips made from GM potatoes (n=489, 43.6%), chicken thighs fed on GM corn (n=472, 42.1%), corn flakes made from GM corn (n=471, 42.0%), GM apples (n=420, 37.4%), wine brewed with GM yeast (416, 37.1%), GM tomatoes (n=408, 36.4%), GM chicken thighs (n=360, 32.1%), GM salmon (n=349, 31.1%).

**Table 2.**
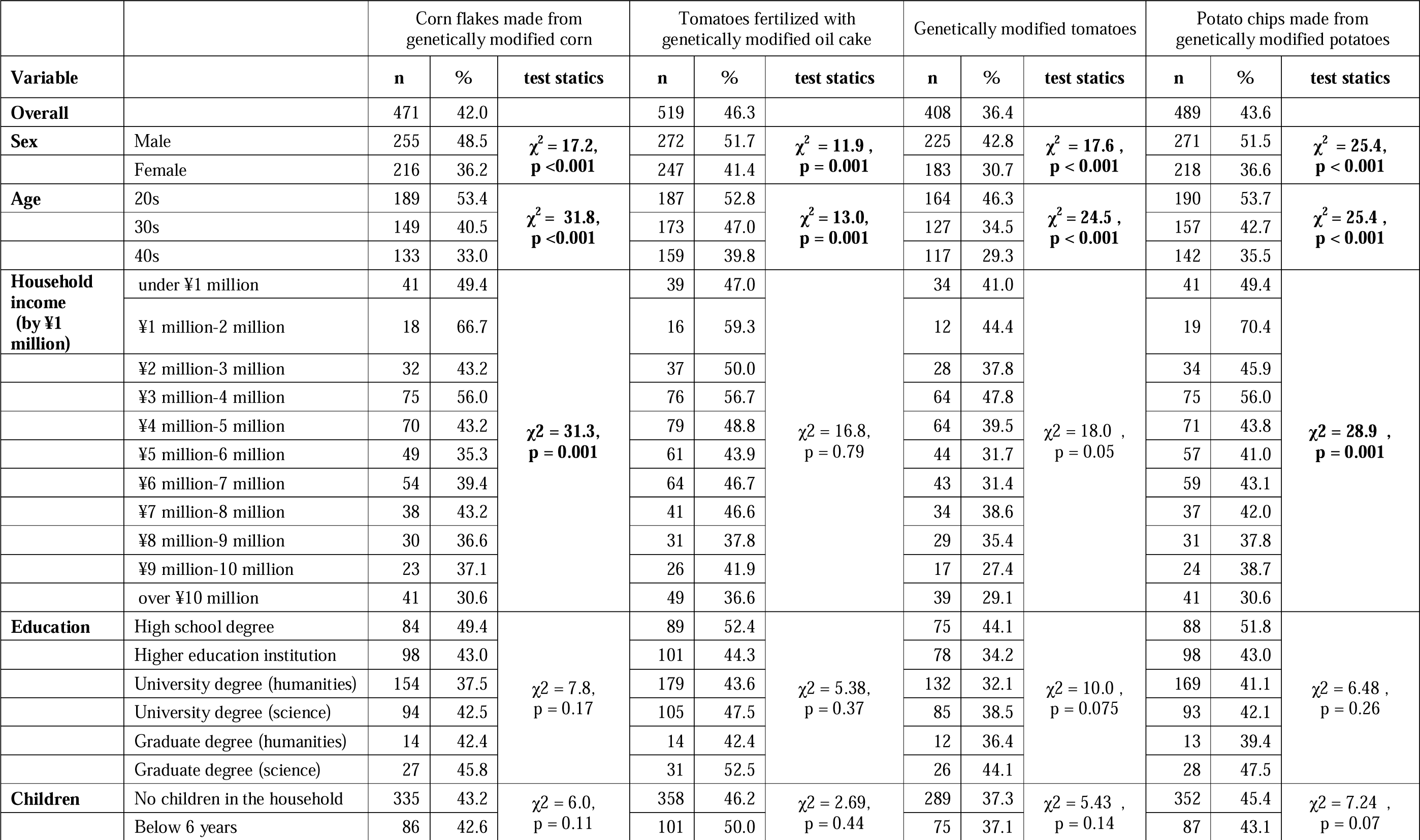
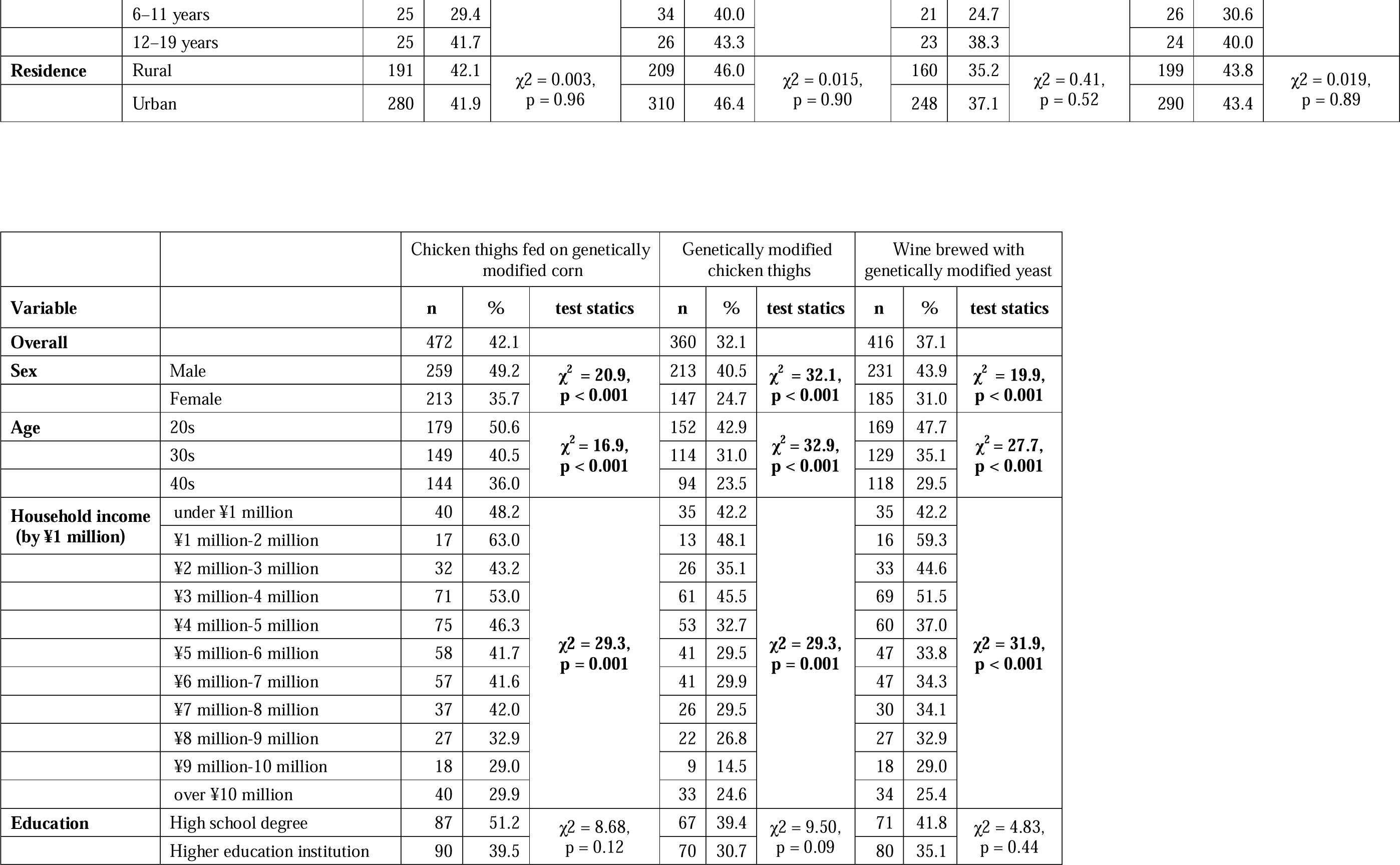
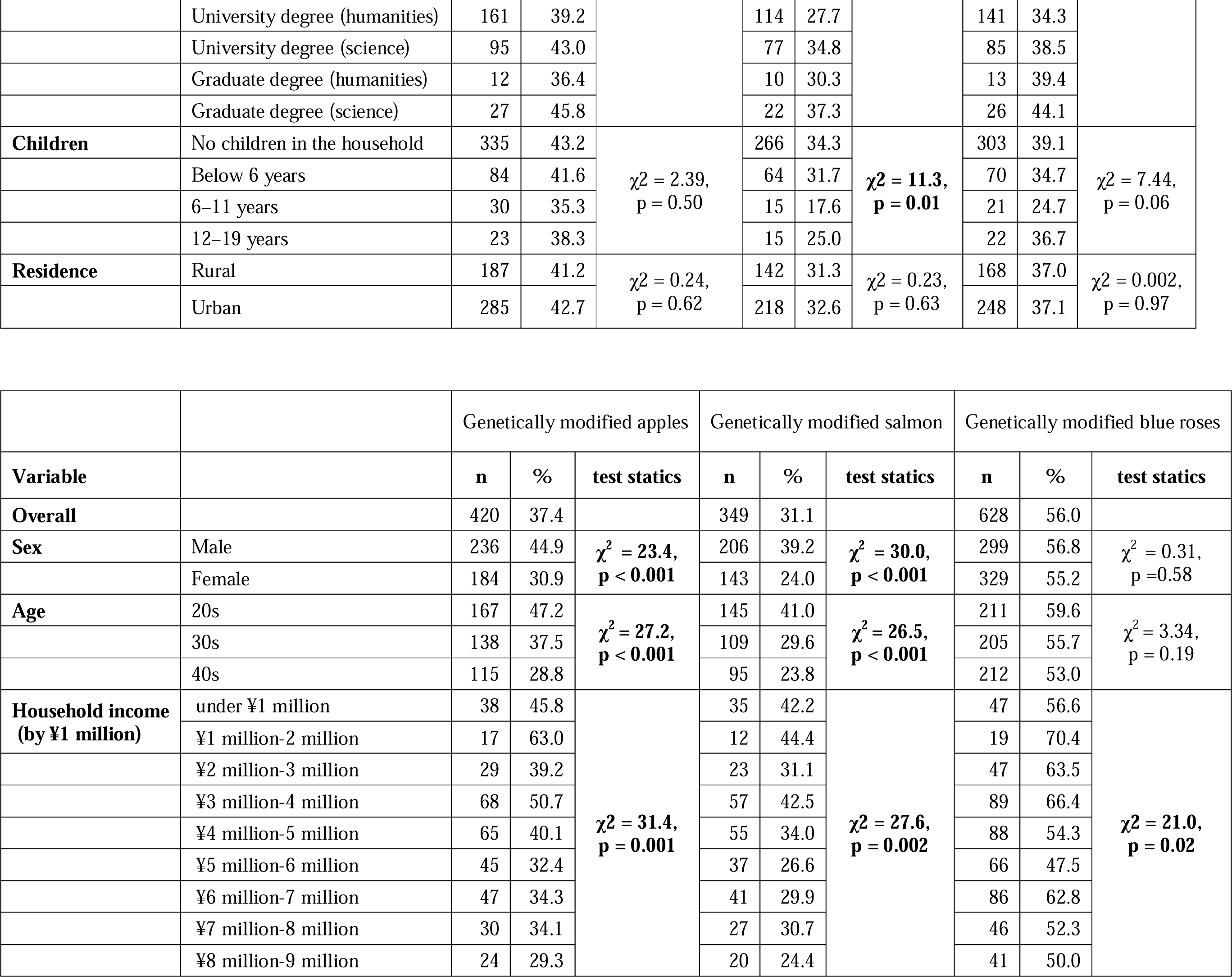
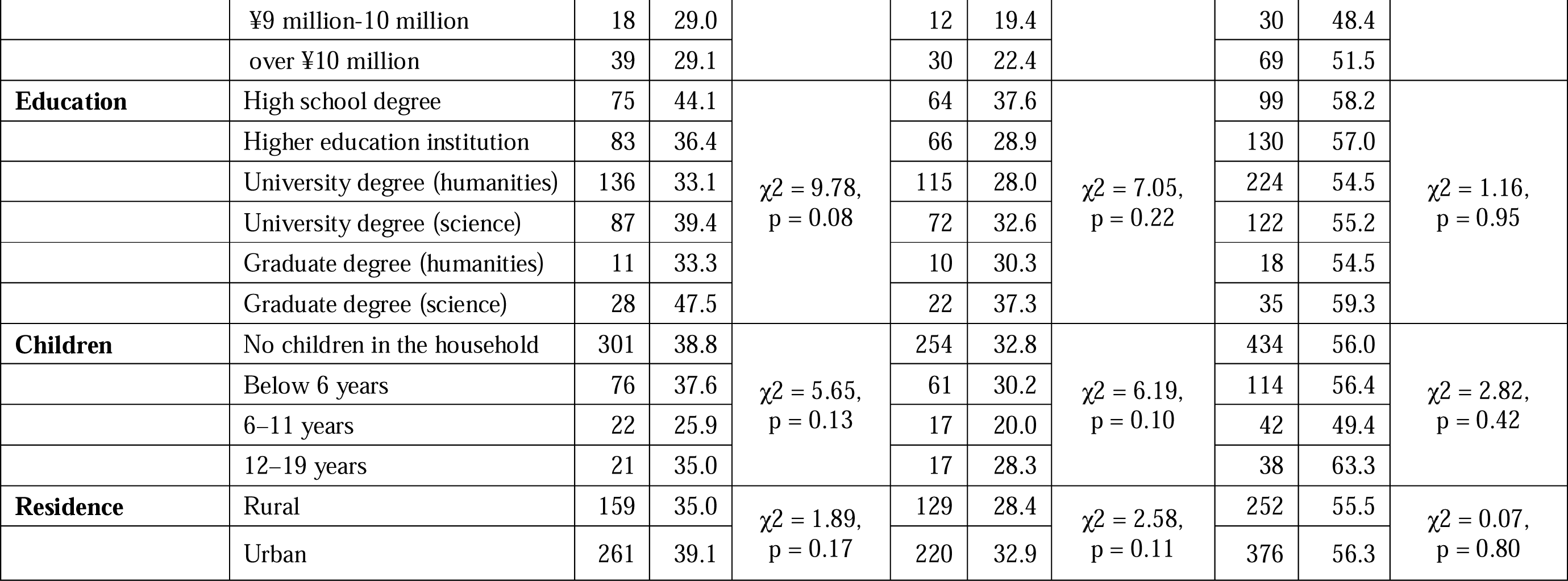
Participants purchase intention (n = 1122).

Willingness to purchase GM blue roses was the highest among various GM products, with more than half of the participants indicating their will to purchase (n = 628; 56%). However, GM animal foods, such as salmon (n = 349; 31.1%) and chicken (n = 360; 32.1%), demonstrated the lowest willingness to purchase.

Respondents showed a relatively higher willingness to purchase for foods indirectly related to GM, that is, those items that used GM fertilizers or ingredients during breeding or cultivation, such as chicken thigh fed on GM corn (n = 472; 42.1%). However, it was low for GM plant foods, such as GM apples (n = 420; 37.4%), GM wine (n = 416; 37.1%), and GM tomatoes (n = 408; 36.4%).

### Differences in willingness to purchase based on sociodemographic characteristics

Table 2 shows the differences in the willingness to purchase based on sociodemographic characteristics. The results showed that sex and age affect the willingness to purchase for all GM products, except ornamental blue roses. Women and older participants had significantly lower willingness to purchase for GM foods than men and younger participants, respectively, (p < 0.001). Participants with higher income showed significantly lower purchase intention of GM foods, except tomatoes fertilized with GM oil cake and GM tomatoes. There were no significant differences in the purchase intention based on education, children, and residence.

### The WTP of GM products and the WTP discount rate from the average market prices of non-GM products corresponding to GM products

#### Comparing the average discount rate of GM products based on applications

##### Analysis of the WTP of those willing to purchase GM products

Table 3 shows the mean WTP and the WTP discount rate from the average market price. Wine brewed with GM yeast had the highest discount rate (36.7%), followed by GM blue roses (30.3%), GM apples (25.3%), corn flakes made from GM corn (24.1%), chicken thighs raised on GM corn (21.5%), GM salmon (19.5%), GM chicken thighs (17.8%), GM tomatoes (17.2%), potato chips made from GM potatoes (16.4%), and tomatoes fertilized with GM oil cake (11.9%). The WTP discount rates of the GM products were not similar to their willingness to purchase.

**Table 3.** WTP of the participants who indicated the willingness to purchase each GM product and the WTP discount rate from the average market price.

| Item |  | Corn flakes made from<br>genetically modified<br>corn<br>(n = 471) |  |  | Tomatoes fertilized with<br>genetically modified oil<br>cake<br>(n = 519) |  |  | Genetically modified<br>tomatoes<br>(n = 408) |  |  |
| --- | --- | --- | --- | --- | --- | --- | --- | --- | --- | --- |
| Price of Non-GM<br>alternative |  | ¥300/200g |  |  | ¥100/1 |  |  | ¥100/1 |  |  |
| Variable |  | Mean<br>WTP<br>(¥) | Disc<br>ount<br>(%) | test<br>static | Mean<br>WTP<br>(¥) | Disc<br>ount<br>(%) | test<br>static | Mean<br>WTP<br>(¥) | Disc<br>ount<br>(%) | test<br>static |
| **Overall** |  | 227.8 | 24.1 |  | 88.1 | 11.9 |  | 82.8 | 17.2 |  |
| **Sex** | Male | 209.7 | 30.1 | t =<br>17.2,<br>p =<br>0.16 | 84.6 | 15.4 | t =<br>1.58,<br>p =<br>0.21 | 78.5 | 21.5 | t =<br>3.33,<br>p =<br>0.07 |
|  | Female | 249.1 | 17.0 |  | 91.9 | 8.1 |  | 88.0 | 12.0 |  |
| **Age** | 20s | 248.8 | 17.1 | F =<br>1.30,<br>p =<br>0.28 | 98.6 | 1.4 | F =<br>3.96,<br>p =<br>0.02 | 86.9 | 13.1 | F =<br>1.68,<br>p =<br>0.19 |
|  | 30s | 224.8 | 25.1 |  | 84.3 | 15.7 |  | 84.2 | 15.8 |  |
|  | 40s | 201.2 | 32.9 |  | 79.7 | 20.3 |  | 75.4 | 24.6 |  |
| **Household income**<br>**(by ¥1 million)** | under ¥1 million | 333.1 | -11.0 | F =<br>0.82,<br>p =<br>0.61 | 101.8 | -1.8 | F =<br>0.61,<br>p =<br>0.81 | 105.7 | -5.7 | F =<br>1.48,<br>p =<br>0.15 |
|  | ¥1 million-2<br>million | 229.4 | 23.5 |  | 115.6 | -15.6 |  | 104.2 | -4.2 |  |
|  | ¥2 million-3<br>million | 208.7 | 30.4 |  | 80.4 | 19.6 |  | 75.2 | 24.8 |  |
|  | ¥3 million-4<br>million | 226.3 | 24.6 |  | 87.8 | 12.2 |  | 79.3 | 20.7 |  |
|  | ¥4 million-5<br>million | 220.4 | 26.5 |  | 86.9 | 13.1 |  | 87.7 | 12.3 |  |
|  | ¥5 million-6<br>million | 222.9 | 25.7 |  | 83.3 | 16.7 |  | 82.9 | 17.1 |  |
|  | ¥6 million-7<br>million | 232.8 | 22.4 |  | 82.3 | 17.7 |  | 84.0 | 16.0 |  |
|  | ¥7 million-8<br>million | 215.0 | 28.3 |  | 86.9 | 13.1 |  | 70.4 | 29.6 |  |
|  | ¥8 million-9<br>million | 199.6 | 33.5 |  | 85.0 | 15.0 |  | 84.5 | 15.5 |  |
|  | ¥9 million-10<br>million | 220.8 | 26.4 |  | 94.6 | 5.4 |  | 79.4 | 20.6 |  |
|  | over ¥10 million | 187.6 | 37.5 |  | 89.1 | 10.9 |  | 68.6 | 31.4 |  |
| **Education** | High school degree | 203.6 | 32.1 | F =<br>0.38,<br>p =<br>0.86 | 91.0 | 9.0 | F =<br>0.28,<br>p =<br>0.92 | 78.8 | 21.2 | F =<br>0.22,<br>p =<br>0.96 |
|  | Higher education<br>institution | 223.1 | 25.6 |  | 84.7 | 15.3 |  | 80.7 | 19.3 |  |
|  | University degree<br>(humanities) | 249.8 | 16.7 |  | 90.8 | 9.2 |  | 85.2 | 14.8 |  |
|  | University degree<br>(science) | 220.2 | 26.6 |  | 87.5 | 12.5 |  | 85.3 | 14.7 |  |
|  | Graduate degree<br>(humanities) | 232.1 | 22.6 |  | 84.3 | 15.7 |  | 80.0 | 20.0 |  |
|  | Graduate degree<br>(science) | 218.5 | 27.2 |  | 78.7 | 21.3 |  | 81.0 | 19.0 |  |
| **Children** | No children in the<br>household | 236.4 | 21.2 | F =<br>0.42,<br>p =<br>0.73 | 90.1 | 9.9 | F =<br>0.89,<br>p =<br>0.45 | 83.8 | 16.2 | F =<br>0.31,<br>p =<br>0.82 |
|  | Below 6 years | 202.9 | 32.4 |  | 81.2 | 18.8 |  | 82.8 | 17.2 |  |
|  | 6–11 years | 214.2 | 28.6 |  | 95.8 | 4.2 |  | 77.5 | 22.5 |  |
|  | 12–19 years | 211.9 | 29.4 |  | 76.5 | 23.5 |  | 74.1 | 25.9 |  |
| **Residence** | Rural | 235.6 | 21.5 | t =<br>0.29,<br>p =<br>0.59 | 88.6 | 11.4 | t =<br>0.02,<br>p =<br>0.88 | 82.8 | 17.2 | t =<br>0.00,<br>p =<br>0.98 |
|  | Urban | 222.4 | 25.9 |  | 87.7 | 12.3 |  | 82.7 | 17.3 |  |

| Item |  | Potato chips made from genetically modified potatoes (n = 489) |  |  | Chicken thighs fed on genetically modified corn (n = 472) |  |  | Genetically modified chicken thighs (n = 360) |  |  |
| --- | --- | --- | --- | --- | --- | --- | --- | --- | --- | --- |
| Price of Non-GM alternative |  | ¥100/60g |  |  | ¥150/100g |  |  | ¥150/100g |  |  |
| Variable |  | Mean WTP (¥) | Discount (%) | test static | Mean WTP (¥) | Discount (%) | test static | Mean WTP (¥) | Discount (%) | test static |
| Overall |  | 83.6 | 16.4 |  | 117.8 | 21.5 |  | 123.3 | 17.8 |  |
| Sex | Male | 82.2 | 17.8 | t = 0.36, p = 0.55 | 115.8 | 22.8 | t = 0.24, p = 0.63 | 120.6 | 19.6 | t = 0.38, p = 0.54 |
|  | Female | 85.4 | 14.6 |  | 120.2 | 19.9 |  | 127.3 | 15.1 |  |
| Age | 20s | 89.5 | 10.5 | F = 3.00, p = 0.05 | 125.0 | 16.7 | F = 2.47, p = 0.09 | 127.2 | 15.2 | F = 0.83, p = 0.53 |
|  | 30s | 85.4 | 14.6 |  | 123.4 | 17.7 |  | 131.8 | 12.1 |  |
|  | 40s | 73.8 | 26.2 |  | 103.0 | 31.3 |  | 106.9 | 28.7 |  |
| Household income (by ¥1 million) | under ¥1 million | 85.7 | 14.3 | F = 0.82, p = 0.61 | 147.3 | 1.8 | F = 1.10, p = 0.36 | 159.5 | -6.3 | F = 1.05, p = 0.40 |
|  | ¥1 million-2 million | 76.1 | 23.9 |  | 103.5 | 31.0 |  | 106.2 | 29.2 |  |
|  | ¥2 million-3 million | 78.9 | 21.1 |  | 115.7 | 22.9 |  | 116.0 | 22.7 |  |
|  | ¥3 million-4 million | 96.9 | 3.1 |  | 111.9 | 25.4 |  | 113.5 | 24.3 |  |
|  | ¥4 million-5 million | 80.7 | 19.3 |  | 109.7 | 26.9 |  | 122.2 | 18.5 |  |
|  | ¥5 million-6 million | 86.2 | 13.8 |  | 141.6 | 5.6 |  | 136.8 | 8.8 |  |
|  | ¥6 million-7 million | 89.5 | 10.5 |  | 120.5 | 19.7 |  | 142.1 | 5.3 |  |
|  | ¥7 million-8 million | 76.8 | 23.2 |  | 111.6 | 25.6 |  | 108.9 | 27.4 |  |
|  | ¥8 million-9 million | 81.0 | 19.0 |  | 116.1 | 22.6 |  | 115.5 | 23.0 |  |
|  | ¥9 million-10 million | 77.0 | 23.0 |  | 103.7 | 30.9 |  | 111.1 | 25.9 |  |
|  | over ¥10 million | 70.0 | 30.0 |  | 96.7 | 35.5 |  | 97.1 | 35.3 |  |
| Education | High school degree | 79.5 | 20.5 | F = 0.52, p = 0.76 | 114.1 | 23.9 | F = 1.14, p = 0.34 | 116.9 | 22.1 | F = 0.12, p = 0.99 |
|  | Higher education institution | 81.5 | 18.5 |  | 108.2 | 27.9 |  | 123.4 | 17.7 |  |
|  | University degree (humanities) | 89.4 | 10.6 |  | 121.1 | 19.3 |  | 128.6 | 14.3 |  |
|  | University degree (science) | 81.5 | 18.5 |  | 113.5 | 24.3 |  | 121.0 | 19.3 |  |
|  | Graduate degree (humanities) | 80.4 | 19.6 |  | 120.0 | 20.0 |  | 127.0 | 15.3 |  |
|  | Graduate degree (science) | 77.5 | 22.5 |  | 156.1 | -4.1 |  | 121.4 | 19.1 |  |
| Children | No children in the household | 85.5 | 14.5 | F = 0.61, p = 0.61 | 121.1 | 19.3 | F = 0.45, p = 0.72 | 124.2 | 17.2 | F = 0.17, p = 0.92 |
|  | Below 6 years | 80.9 | 19.1 |  | 110.4 | 26.4 |  | 125.5 | 16.3 |  |
|  | 6–11 years | 78.3 | 21.7 |  | 109.6 | 26.9 |  | 109.7 | 26.9 |  |
|  | 12–19 years | 71.1 | 28.9 |  | 107.7 | 28.2 |  | 112.0 | 25.3 |  |
| Residence | Rural | 84.6 | 15.4 | t = 0.09, p = 0.76 | 116.6 | 22.3 | t = 0.05, p = 0.83 | 123.5 | 17.7 | t = 0.00, p = 0.98 |
|  | Urban | 82.9 | 17.1 |  | 118.6 | 20.9 |  | 123.2 | 17.9 |  |

| Item |  | Wine brewed with<br>genetically modified<br>yeast<br>(n = 416) |  |  | Genetically modified<br>apples<br>(n = 420) |  |  | Genetically modified<br>salmon<br>(n = 349) |  |  |
| --- | --- | --- | --- | --- | --- | --- | --- | --- | --- | --- |
| Price of Non-GM alternative |  | ¥2000/bottle |  |  | ¥150/1 |  |  | ¥130/1 |  |  |
| Variable |  | Mean<br>WTP (¥) | Disc<br>ount<br>(%) | test<br>static | Mean<br>WTP (¥) | Disc<br>ount<br>(%) | test<br>static | Mean<br>WTP (¥) | Disc<br>ount<br>(%) | test<br>static |
| Overall |  | 1266.0 | 36.7 |  | 112.1 | 25.3 |  | 104.6 | 19.5 |  |
| Sex | Male | 1261.8 | 36.9 | t =<br>0.02,<br>p =<br>0.88 | 111.2 | 25.9 | t =<br>0.06,<br>p =<br>0.81 | 101.1 | 22.2 | t =<br>1.06,<br>p =<br>0.30 |
|  | Female | 1271.1 | 36.4 |  | 113.2 | 24.5 |  | 109.6 | 15.7 |  |
| Age | 20s | 1419.5 | 29.0 | F =<br>9.37,<br>p <<br>0.001 | 118.8 | 20.8 | F =<br>1.16,<br>p =<br>0.31 | 114.0 | 12.3 | F =<br>2.04,<br>p =<br>0.13 |
|  | 30s | 1208.7 | 39.6 |  | 111.3 | 25.8 |  | 99.8 | 23.2 |  |
|  | 40s | 1108.4 | 44.6 |  | 103.4 | 31.1 |  | 95.5 | 26.5 |  |
| Household<br>income<br>(by ¥1 million) | under ¥1 million | 1387.6 | 30.6 | F =<br>2.17,<br>p =<br>0.02 | 140.3 | 6.5 | F =<br>1.14,<br>p =<br>0.33 | 143.4 | -10.3 | F =<br>1.32,<br>p =<br>0.22 |
|  | ¥1 million-2 million | 812.5 | 59.4 |  | 99.4 | 33.7 |  | 89.2 | 31.4 |  |
|  | ¥2 million-3 million | 1368.2 | 31.6 |  | 102.7 | 31.5 |  | 94.4 | 27.4 |  |
|  | ¥3 million-4 million | 1162.5 | 41.9 |  | 100.7 | 32.9 |  | 99.1 | 23.8 |  |
|  | ¥4 million-5 million | 1190.5 | 40.5 |  | 129.2 | 13.9 |  | 104.6 | 19.5 |  |
|  | ¥5 million-6 million | 1231.3 | 38.4 |  | 109.1 | 27.3 |  | 109.2 | 16.0 |  |
|  | ¥6 million-7 million | 1468.9 | 26.6 |  | 116.1 | 22.6 |  | 109.2 | 16.0 |  |
|  | ¥7 million-8 million | 1306.0 | 34.7 |  | 109.5 | 27.0 |  | 94.2 | 27.5 |  |
|  | ¥8 million-9 million | 1338.5 | 33.1 |  | 106.3 | 29.1 |  | 97.0 | 25.4 |  |
|  | ¥9 million-10 million | 1475.0 | 26.3 |  | 112.2 | 25.2 |  | 101.3 | 22.1 |  |
|  | over ¥10 million | 1161.4 | 41.9 |  | 92.7 | 38.2 |  | 87.0 | 33.1 |  |
| Education | High school degree | 1172.9 | 41.4 | F =<br>2.67,<br>p =<br>0.02 | 104.5 | 30.3 | F =<br>1.73,<br>p =<br>0.13 | 95.6 | 26.5 | F =<br>1.18,<br>p =<br>0.32 |
|  | Higher education institution | 1084.8 | 45.8 |  | 92.3 | 38.5 |  | 90.7 | 30.2 |  |
|  | University degree (humanities) | 1318.1 | 34.1 |  | 118.5 | 21.0 |  | 116.2 | 10.6 |  |
|  | University degree (science) | 1376.0 | 31.2 |  | 126.4 | 15.7 |  | 105.7 | 18.7 |  |
|  | Graduate degree (humanities) | 1484.6 | 25.8 |  | 116.4 | 22.4 |  | 110.0 | 15.4 |  |
|  | Graduate degree (science) | 1325.4 | 33.7 |  | 114.0 | 24.0 |  | 105.2 | 19.1 |  |
| Children | No children in the household | 1284.9 | 35.8 | F =<br>0.99,<br>p =<br>0.40 | 113.1 | 24.6 | F =<br>0.14,<br>p =<br>0.93 | 108.1 | 16.8 | F =<br>0.67,<br>p =<br>0.57 |
|  | Below 6 years | 1278.6 | 36.1 |  | 112.6 | 24.9 |  | 95.2 | 26.8 |  |
|  | 6–11 years | 1067.6 | 46.6 |  | 102.3 | 31.8 |  | 95.5 | 26.5 |  |
|  | 12–19 years | 1156.8 | 42.2 |  | 106.7 | 28.9 |  | 94.7 | 27.2 |  |
| Residence | Rural | 1260.1 | 37.0 | t =<br>0.02,<br>p =<br>0.88 | 111.7 | 25.5 | t =<br>0.01,<br>p =<br>0.94 | 107.0 | 17.7 | t =<br>0.21,<br>p =<br>0.65 |
|  | Urban | 1269.9 | 36.5 |  | 112.3 | 25.1 |  | 103.1 | 20.7 |  |

| Item |  | Genetically modified blue roses<br>(n = 628) |
| --- | --- | --- |

| Price of Non-GM alternative |  | ¥800/1 |  |  |
| --- | --- | --- | --- | --- |
| Variable |  | Mean WTP (¥) | Discount(%) | test static |
| **Overall** |  | 557.9 | 30.3 |  |
| **Sex** | Male | 587.2 | 26.6 | t = 1.68,<br>p = 0.20 |
|  | Female | 531.4 | 33.6 |  |
| **Age** | 20s | 575.8 | 28.0 | **F = 3.20,**<br>**p = 0.04** |
|  | 30s | 614.7 | 23.2 |  |
|  | 40s | 485.3 | 39.3 |  |
| **Household income**<br>**(by ¥1 million)** | under ¥1 million | 608.6 | 23.9 | F = 1.02,<br>p = 0.42 |
|  | ¥1 million-2 million | 442.1 | 44.7 |  |
|  | ¥2 million-3 million | 569.2 | 28.9 |  |
|  | ¥3 million-4 million | 525.8 | 34.3 |  |
|  | ¥4 million-5 million | 680.6 | 14.9 |  |
|  | ¥5 million-6 million | 615.8 | 23.0 |  |
|  | ¥6 million-7 million | 551.7 | 31.0 |  |
|  | ¥7 million-8 million | 527.6 | 34.1 |  |
|  | ¥8 million-9 million | 489.0 | 38.9 |  |
|  | ¥9 million-10 million | 530.0 | 33.8 |  |
|  | over ¥10 million | 458.4 | 42.7 |  |
| **Education** | High school degree | 476.8 | 40.4 | F = 1.70,<br>p = 0.13 |
|  | Higher education institution | 510.2 | 36.2 |  |
|  | University degree (humanities) | 553.4 | 30.8 |  |
|  | University degree (science) | 636.0 | 20.5 |  |
|  | Graduate degree (humanities) | 594.4 | 25.7 |  |
|  | Graduate degree (science) | 702.9 | 12.1 |  |
| **Children** | No children in the household | 564.8 | 29.4 | F = 1.06,<br>p = 0.37 |
|  | Below 6 years | 598.4 | 25.2 |  |
|  | 6–11 years | 458.6 | 42.7 |  |
|  | 12–19 years | 468.4 | 41.5 |  |
| **Residence** | Rural | 537.4 | 32.8 | t = 0.61,<br>p = 0.44 |
|  | Urban | 571.7 | 28.5 |  |

##### Analysis of the adjusted WTP

Table 4 presents the mean adjusted WTP and its discount rate from the average market price with WTP = 0 for participants with no purchase intention. Wine made with GM yeast had the highest discount rate (76.5%), followed by GM salmon (75%), GM chicken thighs (73.6%), GM apples (72%), GM tomatoes (69.9%), cornflakes made from GM corn (68.1%), chicken thighs raised on GM corn (67%), potato chips made from GM potatoes (63.6%), GM blue roses (61%), and tomatoes fertilized with GM oil cake (59.3%).

**Table 4.** Adjusted WTP and its discount rate.

| Item |  | Corn flakes made from genetically modified corn<br>(n = 471) |  |  | Tomatoes fertilized with genetically modified oil cake<br>(n = 519) |  |  | Genetically modified tomatoes<br>(n = 408) |  |  |
| --- | --- | --- | --- | --- | --- | --- | --- | --- | --- | --- |
| Price of Non-GM alternative |  | ¥300/200g |  |  | ¥100/1 |  |  | ¥100/1 |  |  |
| Variable |  | Mean WTP (¥) | Discount (%) | test static | Mean WTP (¥) | Discount (%) | test static | Mean WTP (¥) | Discount (%) | test static |
| **Overall** |  | 95.6 | 68.1 |  | 40.7 | 59.3 |  | 30.1 | 69.9 |  |
| **Sex** | Male | 101.7 | 66.1 | t = 0.87,<br>p = 0.35 | 43.7 | 56.3 | t = 1.58,<br>p = 0.21 | 33.6 | 66.4 | **t = 4.60,**<br>**p = 0.03** |
|  | Female | 90.3 | 69.9 |  | 38.1 | 61.9 |  | 27.0 | 73.0 |  |
| **Age** | 20s group | 132.8 | 55.7 | **F = 10.1,**<br>**p < 0.001** | 52.1 | 47.9 | **F = 10.1,**<br>**p < 0.001** | 40.2 | 59.8 | **F = 12.3,**<br>**p < 0.001** |
|  | 30s group | 91.0 | 69.7 |  | 39.6 | 60.4 |  | 29.1 | 70.9 |  |
|  | 40s group | 66.9 | 77.7 |  | 31.7 | 68.3 |  | 22.1 | 77.9 |  |
| **Household income**<br>**(by ¥1 million)** | under ¥1 million | 164.5 | 45.2 | **F = 2.20,**<br>**p = 0.02** | 47.8 | 52.2 | F = 1.39,<br>p = 0.18 | 43.3 | 56.7 | **F = 2.17,**<br>**p = 0.02** |
|  | ¥1 million-2 million | 153.0 | 49.0 |  | 68.5 | 31.5 |  | 46.3 | 53.7 |  |
|  | ¥2 million-3 million | 90.2 | 69.9 |  | 40.2 | 59.8 |  | 28.4 | 71.6 |  |
|  | ¥3 million-4 million | 126.6 | 57.8 |  | 49.8 | 50.2 |  | 37.9 | 62.1 |  |
|  | ¥4 million-5 million | 95.2 | 68.3 |  | 42.4 | 57.6 |  | 34.6 | 65.4 |  |
|  | ¥5 million-6 million | 78.6 | 73.8 |  | 36.5 | 63.5 |  | 26.2 | 73.8 |  |
|  | ¥6 million-7 million | 91.8 | 69.4 |  | 38.4 | 61.6 |  | 26.4 | 73.6 |  |
|  | ¥7 million-8 million | 92.9 | 69.0 |  | 40.5 | 59.5 |  | 27.2 | 72.8 |  |
|  | ¥8 million-9 million | 73.0 | 75.7 |  | 32.1 | 67.9 |  | 29.9 | 70.1 |  |
|  | ¥9 million-10 million | 81.9 | 72.7 |  | 39.7 | 60.3 |  | 21.8 | 78.2 |  |
|  | over ¥10 million | 57.4 | 80.9 |  | 32.6 | 67.4 |  | 20.0 | 80.0 |  |
| **Education** | High school degree | 100.6 | 66.5 | F = 0.04,<br>p = 0.99 | 47.6 | 52.4 | F = 0.61,<br>p = 0.70 | 34.7 | 65.3 | F = 0.90,<br>p = 0.48 |
|  | Higher education institution | 95.9 | 68.0 |  | 37.5 | 62.5 |  | 27.6 | 72.4 |  |
|  | University degree (humanities) | 93.6 | 68.8 |  | 39.5 | 60.5 |  | 27.4 | 72.6 |  |
|  | University degree (science) | 93.7 | 68.8 |  | 41.6 | 58.4 |  | 32.8 | 67.2 |  |
|  | Graduate degree (humanities) | 98.5 | 67.2 |  | 35.8 | 64.2 |  | 29.1 | 70.9 |  |
|  | Graduate degree (science) | 100.0 | 66.7 |  | 41.4 | 58.6 |  | 35.7 | 64.3 |  |
| **Children** | No children in the household | 102.2 | 65.9 | F = 1.16, | 41.6 | 58.4 | F = 0.38, | 31.3 | 68.7 | F = 1.47, |
|  | Below 6 years | 86.4 | 71.2 | p = 0.33 | 40.6 | 59.4 | p = 0.77 | 30.7 | 69.3 | p = 0.22 |
|  | 6–11 years | 63.0 | 79.0 |  | 38.3 | 61.7 |  | 19.2 | 80.8 |  |
|  | 12–19 years | 88.3 | 70.6 |  | 33.1 | 66.9 |  | 28.4 | 71.6 |  |
| **Residence** | Rural | 99.1 | 67.0 | t = 0.23,<br>p = 0.64 | 40.8 | 59.2 | t < 0.01,<br>p = 0.98 | 29.2 | 70.8 | t = 0.24,<br>p = 0.62 |
|  | Urban | 93.2 | 68.9 |  | 40.7 | 59.3 |  | 30.7 | 69.3 |  |

| Item |  | Potato chips made from genetically modified potatoes<br>(n = 489) |  |  | Chicken thighs fed on genetically modified corn<br>(n = 472) |  |  | Genetically modified chicken thighs<br>(n = 360) |  |  |
| --- | --- | --- | --- | --- | --- | --- | --- | --- | --- | --- |
| **Price of Non-GM alternative** |  | ¥100/60g |  |  | ¥150/100g |  |  | ¥150/100g |  |  |
| **Variable** |  | Mean WTP (¥) | Discount (%) | test static | Mean WTP (¥) | Discount (%) | test static | Mean WTP (¥) | Discount (%) | test static |
| **Overall** |  | 36.4 | 63.6 |  | 49.5 | 67.0 |  | 39.6 | 73.6 |  |
| **Sex** | Male | 42.3 | 57.7 | **t = 10.7,**<br>**p = 0.001** | 57.0 | 62.0 | **t = 7.67,**<br>**p = 0.006** | 48.8 | 67.5 | **t = 12.9,**<br>**p < 0.001** |
|  | Female | 31.2 | 68.8 |  | 42.9 | 71.4 |  | 31.4 | 79.1 |  |
| **Age** | 20s group | 48.1 | 51.9 | **F = 14.1,**<br>**p < 0.001** | 63.2 | 57.9 | **F = 8.95,**<br>**p < 0.001** | 54.6 | 63.6 | **F = 12.6,**<br>**p < 0.001** |
|  | 30s group | 36.4 | 63.6 |  | 50.0 | 66.7 |  | 40.8 | 72.8 |  |
|  | 40s group | 26.2 | 73.8 |  | 37.1 | 75.3 |  | 25.1 | 83.3 |  |
| **Household income**<br>**(by ¥1 million)** | under ¥1 million | 42.3 | 57.7 | **F = 2.86,**<br>**p = 0.002** | 71.0 | 52.7 | **F = 2.25,**<br>**p = 0.01** | 67.3 | 55.1 | **F = 2.53,**<br>**p = 0.005** |
|  | ¥1 million-2 million | 53.5 | 46.5 |  | 65.2 | 56.5 |  | 51.1 | 65.9 |  |
|  | ¥2 million-3 million | 36.3 | 63.7 |  | 50.0 | 66.7 |  | 40.8 | 72.8 |  |
|  | ¥3 million-4 million | 54.2 | 45.8 |  | 59.3 | 60.5 |  | 51.7 | 65.5 |  |
|  | ¥4 million-5 million | 35.3 | 64.7 |  | 50.8 | 66.1 |  | 40.0 | 73.3 |  |
|  | ¥5 million-6 million | 35.3 | 64.7 |  | 59.1 | 60.6 |  | 40.4 | 73.1 |  |
|  | ¥6 million-7 million | 38.5 | 61.5 |  | 50.1 | 66.6 |  | 42.5 | 71.7 |  |
|  | ¥7 million-8 million | 32.3 | 67.7 |  | 46.9 | 68.7 |  | 32.2 | 78.5 |  |
|  | ¥8 million-9 million | 30.6 | 69.4 |  | 38.2 | 74.5 |  | 31.0 | 79.3 |  |
|  | ¥9 million-10 million | 29.8 | 70.2 |  | 30.1 | 79.9 |  | 16.1 | 89.3 |  |
|  | over ¥10 million | 21.4 | 78.6 |  | 28.9 | 80.7 |  | 23.9 | 84.1 |  |
| **Education** | High school degree | 41.2 | 58.8 | F = 0.37,<br>p = 0.87 | 58.3 | 61.1 | F = 1.52,<br>p = 0.18 | 46.1 | 69.3 | F = 0.53,<br>p = 0.76 |
|  | Higher education institution | 35.0 | 65.0 |  | 42.7 | 71.5 |  | 37.9 | 74.7 |  |
|  | University degree (humanities) | 36.8 | 63.2 |  | 47.4 | 68.4 |  | 35.7 | 76.2 |  |
|  | University degree (science) | 34.3 | 65.7 |  | 48.8 | 67.5 |  | 42.2 | 71.9 |  |
|  | Graduate degree (humanities) | 31.7 | 68.3 |  | 43.6 | 70.9 |  | 38.5 | 74.3 |  |
|  | Graduate degree (science) | 36.8 | 63.2 |  | 71.4 | 52.4 |  | 45.3 | 69.8 |  |
| **Children** | No children in the household | 38.8 | 61.2 | F = 2.28,<br>p = 0.08 | 52.3 | 65.1 | F = 1.05,<br>p = 0.37 | 42.6 | 71.6 | F = 2.51,<br>p = 0.06 |
|  | Below 6 years | 34.9 | 65.1 |  | 45.9 | 69.4 |  | 39.7 | 73.5 |  |
|  | 6–11 years | 24.0 | 76.0 |  | 38.7 | 74.2 |  | 19.4 | 87.1 |  |
|  | 12–19 years | 28.5 | 71.5 |  | 41.3 | 72.5 |  | 28.0 | 81.3 |  |
| **Residence** | Rural | 37.1 | 62.9 | t = 0.10,<br>p = 0.76 | 48.0 | 68.0 | t = 0.25,<br>p = 0.62 | 38.6 | 74.3 | t = 0.10,<br>p = 0.75 |
|  | Urban | 36.0 | 64.0 |  | 50.6 | 66.3 |  | 40.2 | 73.2 |  |

| Item |  | Wine brewed with genetically modified yeast<br>(n = 416) |  |  | Genetically modified apples<br>(n = 420) |  |  | Genetically modified salmon<br>(n = 349) |  |  |
| --- | --- | --- | --- | --- | --- | --- | --- | --- | --- | --- |
| **Price of Non-GM alternative** |  | ¥2000/bottle |  |  | ¥150/1 |  |  | ¥130/1 |  |  |
| **Variable** |  | Mean WTP (¥) | Discount (%) | test static | Mean WTP (¥) | Discount (%) | test static | Mean WTP (¥) | Discount (%) | test static |
| **Overall** |  | 469.4 | 76.5 |  | 42.0 | 72.0 |  | 32.5 | 75.0 |  |
| **Sex** | Male | 554.1 | 72.3 | t = 13.7,<br>p < 0.001 | 49.9 | 66.7 | t = 11.3,<br>p = 0.001 | 39.6 | 69.5 | t = 12.1,<br>p = 0.001 |
|  | Female | 394.6 | 80.3 |  | 34.9 | 76.7 |  | 26.3 | 79.8 |  |
| **Age** | 20s group | 677.7 | 66.1 | F = 24.04,<br>p < 0.001 | 56.0 | 62.7 | F = 11.9,<br>p < 0.001 | 46.7 | 64.1 | F = 14.0,<br>p < 0.001 |
|  | 30s group | 423.7 | 78.8 |  | 41.7 | 72.2 |  | 29.6 | 77.2 |  |
|  | 40s group | 327.0 | 83.7 |  | 29.7 | 80.2 |  | 22.7 | 82.5 |  |
| **Household income**<br>**(by ¥1 million)** | under ¥1 million | 585.1 | 70.7 | F = 1.89,<br>p = 0.04 | 64.2 | 57.2 | F = 2.42,<br>p = 0.008 | 60.5 | 53.5 | F = 3.04,<br>p = 0.001 |
|  | ¥1 million-2 million | 481.5 | 75.9 |  | 62.6 | 58.3 |  | 39.6 | 69.5 |  |
|  | ¥2 million-3 million | 610.1 | 69.5 |  | 40.2 | 73.2 |  | 29.3 | 77.5 |  |
|  | ¥3 million-4 million | 598.6 | 70.1 |  | 51.1 | 65.9 |  | 42.1 | 67.6 |  |
|  | ¥4 million-5 million | 440.9 | 78.0 |  | 51.8 | 65.5 |  | 35.5 | 72.7 |  |
|  | ¥5 million-6 million | 416.3 | 79.2 |  | 35.3 | 76.5 |  | 29.1 | 77.6 |  |
|  | ¥6 million-7 million | 503.9 | 74.8 |  | 39.8 | 73.5 |  | 32.7 | 74.8 |  |
|  | ¥7 million-8 million | 445.2 | 77.7 |  | 37.3 | 75.1 |  | 28.9 | 77.8 |  |
|  | ¥8 million-9 million | 440.7 | 78.0 |  | 31.1 | 79.3 |  | 23.7 | 81.8 |  |
|  | ¥9 million-10 million | 428.2 | 78.6 |  | 32.6 | 78.3 |  | 19.6 | 84.9 |  |
|  | over ¥10 million | 294.7 | 85.3 |  | 27.0 | 82.0 |  | 19.5 | 85.0 |  |
| **Education** | High school degree | 489.9 | 75.5 | **F = 1.53,**<br>**p = 0.18** | 46.1 | 69.3 | **F = 1.59,**<br>**p = 0.16** | 36.0 | 72.3 | **F = 0.70,**<br>**p = 0.62** |
|  | Higher education institution | 380.6 | 81.0 |  | 33.6 | 77.6 |  | 26.3 | 79.8 |  |
|  | University degree (humanities) | 452.2 | 77.4 |  | 39.2 | 73.9 |  | 32.5 | 75.0 |  |
|  | University degree (science) | 529.2 | 73.5 |  | 49.8 | 66.8 |  | 34.4 | 73.5 |  |
|  | Graduate degree (humanities) | 584.8 | 70.8 |  | 38.8 | 74.1 |  | 33.3 | 74.4 |  |
|  | Graduate degree (science) | 584.1 | 70.8 |  | 54.1 | 63.9 |  | 39.2 | 69.8 |  |
| **Children** | No children in the household | 502.4 | 74.9 | **F = 3.00,**<br>**p = 0.03** | 43.9 | 70.7 | **F = 1.47,**<br>**p = 0.22** | 35.4 | 72.8 | **F = 2.16,**<br>**p = 0.09** |
|  | Below 6 years | 442.7 | 77.9 |  | 42.4 | 71.7 |  | 28.8 | 77.8 |  |
|  | 6–11 years | 263.8 | 86.8 |  | 26.5 | 82.3 |  | 19.1 | 85.3 |  |
|  | 12–19 years | 424.2 | 78.8 |  | 37.3 | 75.1 |  | 26.8 | 79.4 |  |
| **Residence** | Rural | 466.3 | 76.7 | **t = 0.01,**<br>**p = 0.91** | 39.1 | 73.9 | **t = 1.10,**<br>**p = 0.30** | 30.4 | 76.6 | **t = 0.84,**<br>**p = 0.36** |
|  | Urban | 471.5 | 76.4 |  | 43.9 | 70.7 |  | 34.0 | 73.8 |  |

| Item |  | Genetically modified blue roses<br>(n = 628) |  |  |
| --- | --- | --- | --- | --- |
| **Price of Non-GM alternative** |  | ¥800/1 |  |  |
| **Variable** |  | Mean WTP (¥) | Discount(%) | test static |
| **Overall** |  | 312.3 | 61.0 |  |
| **Sex** | Male | 333.8 | 58.3 | **t = 1.91,**<br>**p = 0.17** |
|  | Female | 293.3 | 63.3 |  |
| **Age** | 20s group | 343.2 | 57.1 | **F = 3.96,**<br>**p = 0.02** |
|  | 30s group | 342.4 | 57.2 |  |
|  | 40s group | 257.2 | 67.9 |  |
| **Household income**<br>**(by ¥1 million)** | under ¥1 million | 344.6 | 56.9 | **F = 1.11,**<br>**p = 0.35** |
|  | ¥1 million-2 million | 311.1 | 61.1 |  |
|  | ¥2 million-3 million | 361.5 | 54.8 |  |
|  | ¥3 million-4 million | 349.2 | 56.4 |  |
|  | ¥4 million-5 million | 369.7 | 53.8 |  |
|  | ¥5 million-6 million | 292.4 | 63.5 |  |
|  | ¥6 million-7 million | 346.3 | 56.7 |  |
|  | ¥7 million-8 million | 275.8 | 65.5 |  |
|  | ¥8 million-9 million | 244.5 | 69.4 |  |
|  | ¥9 million-10 million | 256.6 | 67.9 |  |
|  | over ¥10 million | 236.0 | 70.5 |  |
| **Education** | High school degree | 277.7 | 65.3 | F = 1.12,<br>p = 0.35 |
|  | Higher education institution | 290.9 | 63.6 |  |
|  | University degree (humanities) | 301.6 | 62.3 |  |
|  | University degree (science) | 351.1 | 56.1 |  |
|  | Graduate degree (humanities) | 324.2 | 59.5 |  |
|  | Graduate degree (science) | 416.9 | 47.9 |  |
| **Children** | No children in the household | 316.3 | 60.5 | F = 1.09,<br>p = 0.35 |
|  | Below 6 years | 337.7 | 57.8 |  |
|  | 6–11 years | 226.6 | 71.7 |  |
|  | 12–19 years | 296.7 | 62.9 |  |
| **Residence** | Rural | 298.3 | 62.7 | t = 0.62,<br>p = 0.43 |
|  | Urban | 321.8 | 59.8 |  |

##### Comparing the average discount rate of GM products based on sociodemographic characteristics

Regarding the factors affecting the mean WTP, as shown in Table 3, as participants’ ages increased, the WTP discount rate of GM tomato, GM wine, and GM roses increased. There was a significant difference in the mean WTP of GM wine based on education level. Regarding the factor affecting the mean adjusted WTP of various GM products, as shown in Table 4, there were significant differences in the mean adjusted WTP of all the GM products based on age. Moreover, there were significant differences in the mean adjusted WTP of the majority of GM products, except for corn flakes made from GM corn and tomatoes fertilized with GM oil cake, based on sex.

## Discussion

This study examined the purchase intention for 10 GM-related items, including animal and plant GM foods and ornamental GM roses. These can be divided into five groups – animal products (GM chicken thighs and GM salmon), plant products (GM cornflakes, GM tomatoes, GM potato chips, GM yeast wine, and GM apples), GM-fed plant products (GM oil cake fertilized tomatoes), GM-fed animal products (chicken thighs fed with GM corn), and GM roses for ornamental use. Therefore, we discuss the differences in purchase intention and WTP between food and non-food, plant food and animal food, and plant food and GM-fed animal food.

### Food vs. non-food products

Ornamental roses had the highest percentage of willingness to purchase, the highest WTP discount rate, and the lowest adjusted discount rate. Since the willingness to purchase is an estimate of acceptance, and ornamental roses are not food, these outcomes are consistent with the expectation. Concerns about GM products include human health effects, environmental effects, and ethics [9,13]. This is because the direct impact on the human body is most important for acceptance, even though ornamental roses are considered to have an environmental impact. The high discount rate for GM roses was surprising; however, it may be because the WTP was measured only among those who expressed a purchase intention. Thus, it did not indicate a low level of acceptance in the aggregate. A drawback of WTP is that it is a stated value and outlier values can be large [14]. As shown in S1 Dataset, some respondents stated outliers that were of different orders of magnitude. The analysis was performed without excluding these outliers. This suggests the need to consider how to handle outliers in the WTP. Moreover, among the respondents with an intermediate acceptance level, those who indicated a purchase intention but rated lower WTP were more likely to be included when the purchase intention rate was higher. This would have reduced the mean WTP. Therefore, we calculated the adjusted WTP, with WTP = 0 for those with no purchase intention. This indicated the lowest discount rate for roses, which was consistent with the expectation. However, the high WTP shows the price paid by those with purchase intentions, which is important in the marketplace.

### GM plant food vs GM animal food

In this study, the purchase intention was around 30% for animal foods (GM chicken thighs and GM salmon) and around 40% for plant foods (GM cornflakes, GM tomatoes, GM potato chips, GM yeast wine, and GM apples); a difference of around 10%. Previous studies have shown that the acceptance of animal products is lower than that of plant products [12,13,15]; thus, the results are consistent with those findings. This may be related to the psychological mechanism, whereby, people are ethically resistant to animal products than plant products [13,16].

The WTP discount rates for animal products were around 20% (GM chicken thighs = 17.8% and GM salmon = 19.5%), while it was 15-35% for plant foods (GM corn flakes = 24.1%, GM tomatoes = 17.2%, GM potato chips = 16.4%, GM yeast wine = 36.7%, GM apples = 25.3%). Some GM plants had higher discount rates. This may be due to the limitations of the WTP measurement and the problems with outlier averages (discussed above). The high discount rate for wine may be due to the high original price, similar to the high discount rate for ornamental roses (30%).

Regarding adjusted WTP, the discount rates for animal products were around 70% (GM chicken thighs = 73.6%, GM salmon = 75%), while it was 60-75% for plant foods (GM corn flakes = 68.1%, GM tomatoes = 69.9%, GM potato chips = 63.6%, GM yeast wine = 76.5%, GM apples = 72%). The discount rates were relatively higher for animal products than for plant products. As in the survey by Chern et al., where the Japanese gave a premium of around 30% for non-GM plant foods and a premium of 50-60% for non-GM animal products [17], there were certain differences in the WTP premium in quantity, especially for plant foods. However, it can be inferred that Japanese people have a low acceptance level of GM plant foods, although there is a limitation in the WTP adjustment method. The WTP discount rate for GM plant foods was around 16-37%, which may be helpful from the market’s perspective because GM plant foods are on the market in the form of raw materials, such as oil.

### GM plant foods vs GM-fed animal foods

The purchase intention rates for plant foods using GM feed were as follows – tomatoes fertilized with GM oil cake = 46.3%, GM tomatoes = 36.4%, while for animal products using GM feed, they were as follows – chicken thighs fed GM corn = 42.1%, GM chicken thighs = 32.1%. For both plant and animal products, the purchase intention percentage of products that used GM fertilizers was about 10% higher than those that were directly involved GM. This is because people feel that products with GM fertilizers are indirectly affected by GM technology.

The WTP discount rate difference between GM-fed foods and GM foods were as follows – tomatoes fertilized with GM oil cake = 11.9% and GM tomatoes = 17.2%, and GM-fed animal products = 21.5% and chicken thighs fed with GM corn = 17.8%. The latter results regarding the comparison between GM-fed and GM animals contradict the comparison between GM-fed and GM plants. We consider this to be a limitation of the WTP measurement (discussed above). Therefore, using the adjusted WTP, the discount rate for plant foods using GM feed was as follows – tomatoes fertilized with GM oil cake = 59.3% compared with GM tomatoes = 69.9%, while for animal foods it was as follows – chicken thighs fed with GM corn = 67% compared with GM chicken thighs = 73.6%. Hence, it can be inferred that the discount rate is about 10% lower than GM-fed foods due to the perceived risk of GM products.

We consider the effect of sociodemographic characteristics on the willingness to purchase percentage and WTP. Our study suggests that sex, age, and income are associated with purchase intention and the WTP of GM products.

Based on sex, the percentage of willingness to purchase was significantly lower among women for all GM food products. This is consistent with the results of previous studies [5,18–21]. The purchase intention percentage among men was about 40-50%, whereas it was about 30-40% for women, with differences of about 10% for all items. There were significant differences in the WTP for all GM products between men and women; however, women had a significantly higher discount rate for the adjusted WTP. Although it is difficult to determine why women had less purchase intention than men, it may be because they feel more socially responsible for food safety in their households; thus, have a greater risk perception [22].

Regarding age, as age increased, the percentages of willingness to purchase decreased for all GM foods. Previous studies have not clarified the effect of age on acceptance. Some studies found that older people were less aware of the risks and more accepting [21,23], while others found younger people to be more accepting [24]. The willingness to purchase percentage of people in their 20s was around 40-50%, while it was around 30-40% for people in their 30s. In the WTP, there were significant differences among tomatoes fertilized with GM oil cake and GM wine among different ages. The discount rates for GM-fed tomatoes were 1.4%, 15.7%, and 20.3% for people in their 20s, 30s, and 40s, respectively, and for GM wine were 29%, 39.6%, and 44.6% for people in their 20s, 30s, and 40s, respectively. The discount rate increased significantly with age. For adjusted WTP, the discount rates for GM chicken, salmon, and wine were 60%, 70%, and 80% for respondents in their 20s, 30s, and 40s, respectively. The discount rates increased by about 10% for each age group. This may be related to the social experiences of each age group, especially in terms of family roles, marriage, and childbearing, and the different timing and duration of exposure, since people in their 40s were adults when GM foods first appeared, while those in their 20s were children [7].

Regarding income, the willingness to purchase percentage was lower for people with higher incomes. The purchase intentions were 40–50% for households with annual income below ¥1 million compared to 20–30% for households with annual income above ¥10 million. In addition, although there were no significant differences in the WTP discount rates in this study, in the case of chicken thigh meat, those with annual income below ¥1 million placed a 6.3% premium for GM thigh, while those with annual incomes of more than ¥10 million discounted the GM meat by 35.3%. Previous research has shown that high income is associated with high WTP for GM foods with a variety of benefits [25]. Furthermore, it has been said in the field of psychology that higher-income groups have a lower risk perception for various hazards [22]. However, this study’s results were contrary to those of previous studies. This may have been influenced by the fact that the WTP was asked without highlighting the benefits of GM foods (less pesticides, higher nutritional value, etc.).

Moreover, the higher the income, the more risk-averse people become because they have more choices for purchasing foods.

This study has certain limitations. It has a selection bias because the survey was conducted on the Internet. Secondary data was used for analysis. Furthermore, psychological and social structures that lead to acceptance are complex, and confounding factors need to be included in the statistics; however, the factors examined in this study were limited. The validity of the WTP scale and the possibility of extreme values may distort the mean.

Despite these limitations, this study is meaningful because it is the first study to clarify how Japanese people evaluate and quantitatively compare purchase intention to a wide range of GM products.

## Conclusion

Regarding willingness to purchase various GM products in Japan, the results quantitatively showed that GM animal foods were less accepted than other products, while GM plant foods and ornamental products had high acceptance. The WTP comparison, based on sociodemographic characteristics, suggested that the WTP may be lower for women, older people, and people with higher incomes. These findings are useful for Japan where it is necessary to understand consumer preferences to have good risk communication with consumers. These may serve as a marketing reference for countries worldwide that are facing similar problems.

## Acknowledgments

The authors would like to thank the staff of the Department of Public Health, Health Management and Policy, and Nara Medical University.

## Supporting information

S1 Dataset.

## References

1. ISAAA. Global status of commercialized biotech/GM crops in 2017: Biotech crop adoption surges as economic benefits accumulate in 22 Years. ISAAA Brief No. 53. ISAAA: Ithaca, NY; 2017.

2. Lucht JM. Public acceptance of plant biotechnology and GM crops. Viruses. 2015;7(8): 4254–81.

3. Baumblatt JAG, Carpenter LR, Wiedeman C, Dunn JR, Schaffner W, Jones TF. Population survey of attitudes and beliefs regarding organic, genetically modified, and irradiated foods. Nutr Health. 2017;23(1): 7–11.

4. Komoto K, Okamoto S, Hamada M, Obana N, Samori M, Imamura T. Japanese consumer perceptions of genetically modified food: Findings from an international comparative study. Interact J Med Res. 2016;5:e23.

5. Hoban TJ. Consumer acceptance of biotechnology in the United States and Japan. Food Technol. 1999;53:50–3.

6. Hoban TJ. Consumer acceptance of biotechnology: An international perspective. Nat Biotechnol. 1997;15:232–4.

7. Mine A, Okamoto S, Myojin T, Hamada M, Imamura T. Basic biology education in high school and acceptance of genetically modified food in Japan. PLoS ONE. 2023;18:e0281493.

8. Ministry of Agriculture, Forestry and Fisheries. Domestic and international situation concerning genetically modified agricultural crops (in Japanese). [cited 20 Sep 2023]. Available: https://www.maff.go.jp/j/syouan/nouan/carta/zyoukyou/.

9. Frewer L, Lassen J, Kettlitz B, Scholderer J, Beekman V, Berdal KG. Societal aspects of genetically modified foods. Food Chem Toxicol. 2004;42:1181–93.

10. Venkatachalam L. The contingent valuation method: A review. Environ Impact Assess Rev. 2004;24(1):89–124.

11. Noussair C, Robin S, Ruffieux B. Do consumers really refuse to buy genetically modified food? Econ J. 2004;114:102–20.

12. Chiaravutthi Y, Pongjit C. Thai consumer willingness to pay for differing GM labeling policies: Comparisons across time. Thail World Econ. 2022;40: 69-86.

13. Knight A. Intervening effects of knowledge, morality, trust, and benefits on support for animal and plant biotechnology applications. Risk Anal. 2007;27(6):1553–63.

14. Borzykowski N, Baranzini A, Maradan D. Scope effects in contingent valuation: Does the assumed statistical distribution of WTP matter? Ecol Econ. 2018;144:319–29.

15. Onyango B, Govindasamy R, Hallman WK, Jang H-M, Puduri VS. Consumer acceptance of genetically modified foods in KorealJ: Factor and cluster analysis. J Agribus. 2006;24(1):1–18.

16. Einsiedel E. Cloning and its discontents – a Canadian perspective. Nat Biotechnol. 2000;18:943–4.

17. Chern WS, Rickertsen K, Tsuboi N, Fu T-T. Consumer acceptance and willingness to pay for genetically modified vegetable oil and salmon: A multiple-country assessment. AgBioForum. 2002;5(3):105–12.

18. Ganiere P, Chern WS, Hahn D. A continuum of consumer attitudes toward genetically modified foods in the United States. J Agric Resour Econ. 2006;31:129–49.

19. Hossain F, Onyango B. Product attributes and consumer acceptance of nutritionally enhanced genetically modified foods. Int J Consum Stud. 2004;28:255–67.

20. Siegrist M. Perception of gene technology, and food risks: results of a survey in Switzerland. J Risk Res. 2003;6:45–60.

21. Christoph IB, Bruhn M, Roosen J. Knowledge, attitudes towards and acceptability of genetic modification in Germany. Appetite. 2008;51:58–68.

22. Slovic P. Trust, emotion, sex, politics, and science: Surveying the risk-assessment battlefield. Risk Anal. 1999;19(4):689–701.

23. Popoola LA, Otitoju MA, Tanko L. Determinants of willingness to consume genetically modified (GM) foods and the potential expenditure in Nigeria. GM Crops Food. 2017;12(1):85.

24. Pardo R, Midden C, Miller JD. Attitudes toward biotechnology in the European Union. J Biotechnol. 2002;98:9–24.

25. Boccaletti S, Moro D. Consumer willingness-to-pay for GM food products in Italy.

